# Complexity of dendritic SER increases at enlarging synapses during LTP

**DOI:** 10.1101/015974

**Authors:** Michael A. Chirillo, Jennifer N. Bourne, Laurence F. Lindsey, Kristen M. Harris

## Abstract

Smooth endoplasmic reticulum (SER) forms a membranous network that extends throughout neurons. SER regulates intracellular calcium and the posttranslational modification and trafficking of membrane and proteins. As the structure of dendritic SER shifts from a tubular to a more complex, branched form, the movement of membrane cargo slows and delivery to nearby spines increases. Here we discovered changes in the structural complexity of SER that have important functional implications during long-term potentiation (LTP) in adult rat hippocampus. By 2 hours after the induction of LTP with theta-burst stimulation, synapse enlargement was greatest on spines that contained SER. More spines had an elaborate spine apparatus than a simple tubule of SER. The SER in dendritic shafts became more complex beneath spines with both polyribosomes and SER, and less complex along aspiny dendritic regions. The findings suggest that local changes in dendritic SER support enhanced growth of specific synapses during LTP.

## Introduction

Long-term potentiation (LTP) is a cellular correlate of learning and memory that induces structural plasticity of dendritic spines and synapses^1-3^. To maintain enhanced synaptic transmission during LTP, postsynaptic densities (PSDs) enlarge^4, 5^, endosomal compartments are rapidly engaged^6, 7^, and AMPA receptors are inserted^8, 9^. Polyribosomes, the cell’s protein synthetic machinery, occur in 6-12% of dendritic spines in mature hippocampus, a frequency that fluctuates during different stages of LTP and development^4, 10, 11^. These observations have led to the hypothesis that local sources of membrane and protein synthesis can be mobilized following the induction of LTP and contribute to synapse growth.

Smooth endoplasmic reticulum (SER) is the cell’s largest organelle, extending as a membranous network throughout the processes of neurons, yet we know little about its role during LTP. SER regulates calcium locally and provides posttranslational modification and trafficking of integral membrane proteins^6, 12^. In the dendritic shaft, SER gives rise to local areas of complexity that retain and enhance delivery of cargo to nearby synapses^13^. SER enters less than 20% of CA1 dendritic spines^14^ and can form simple tubules or a spine apparatus, which comprises folds of SER stacked between dense staining plates that contain the actin binding protein synaptopodin^14-16^. The presence of SER in a dendritic spine enhances the local store of calcium^17^ that is released during plasticity-inducing stimuli^18-20^.

In our prior work, we found that a subset of small dendritic spines were eliminated while remaining synapses were enlarged by 2 hours during LTP^4^. Here we hypothesized that redistribution of SER along dendrites and into dendritic spines would determine specifically which synapses would be enlarged during LTP. To test this hypothesis, we analyzed three-dimensional reconstructions from serial section electron microscopy of SER in mature hippocampal CA1 dendrites that had undergone LTP induced with theta-burst stimulation. Four novel and functionally important changes in the structure of SER were discovered. First, SER in dendritic spines was more likely to form a spine apparatus and occupied a greater volume during LTP. Second, SER in dendritic shafts was less complex in regions of the dendrite lacking spines, suggesting a more rapid movement of membrane cargo along regions lacking spines and synapses. Third, the synapses on spines that contained SER were larger in both control and LTP conditions and underwent the most growth during LTP. Finally, during LTP the complexity of SER in the dendritic shaft was conserved beneath most spines and became significantly more complex at the base of spines that contained both polyribosomes and SER. These findings suggest that SER was preferentially redistributed along the dendritic shaft to target membrane trafficking into dendritic spines where synapse growth was greatest and to support local protein synthesis during LTP.

## Results

Slices were prepared from the middle of adult rat hippocampus, and two stimulating electrodes were placed on either side of a recording electrode in the middle of *stratum radiatum* of area CA1 (Fig 1a)^4^. LTP was induced with theta-burst stimulation (TBS) at one stimulating electrode while the other received control pulses. At 2 hours post-TBS, the slices were rapidly fixed, processed, and prepared for 3DEM (Fig. 1b, see Methods). SER was identified on the basis of its appearance as irregularly shaped, membranous cisternae with a clear lumen (Fig. 1c)^13, 14^.

**Figure 1:**
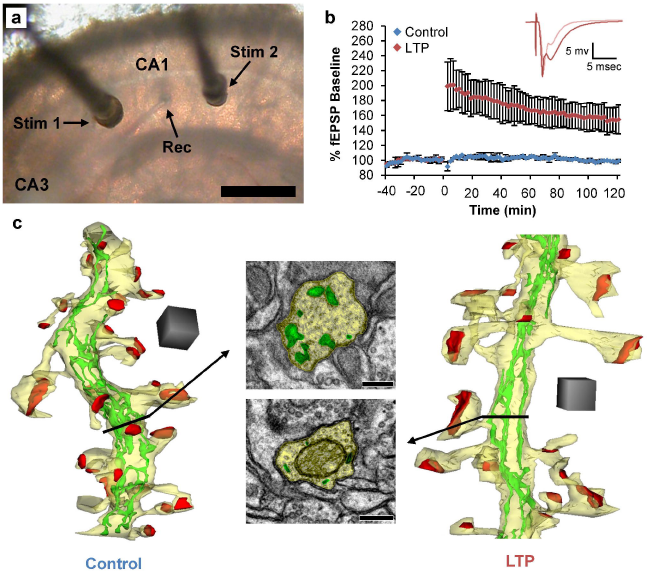
LTP induction and SER identification in hippocampal area CA1 of adult rats. (a) Two stimulating electrodes (Stim 1 and 2) were positioned 600-800 μm apart on either side of a recording electrode (Rec) in the middle of *stratum radiatum* in hippocampal area CA1. Scale bar = 600 μm. (b) Baseline responses were collected for ~40 min. TBS was delivered to one stimulating electrode to induce LTP while the other received control stimulation. Postsynaptic responses were monitored for 2 hours post-TBS, after which the tissue was rapidly fixed (within 30 sec of last test pulse) and prepared for EM. Inset shows example waveforms from responses to the stimulating electrode where LTP was induced, before (pink) and after (red) receiving TBS. (c) Example reconstructions of dendrites (yellow) with associated PSDs (red) and SER (green) in the dendritic shaft. Scale cubes = 0.5 pm on each side. Two micrographs illustrate a section of the dendritic shaft (yellow) from the dendrites with SER (green). SER was identified on EM sections as membranous cisternae with a clear lumen. Scale bar = 250 μm.

In control conditions, we found that the volume of SER per unit length of dendrite was strongly and positively correlated with the amount of synaptic input supported by the dendritic segment (Fig. 2). Interestingly, even though the total amount of SER per length of dendrite did not change significantly during LTP, the correlation between SER volume and synaptic input broke down by 2 hours during LTP, suggesting that SER was being remodeled. This finding prompted us to investigate the underlying structural alterations in SER that might be occurring during plasticity.

**Fig. 2:**
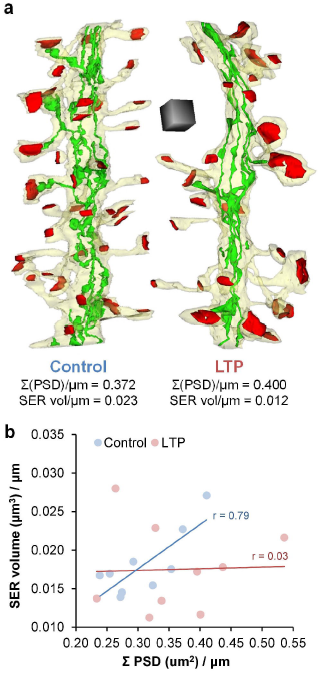
Correlation between total synaptic input and SER volume along the dendrite breaks down by 2 hours during LTP. (a) Examples of reconstructed dendritic segments (yellow) with synapses (red) and SER (green). Scale cube = 0.5 μm on each side. (b) SER volume versus summed PSD area per unit length of dendrite in both conditions. Total SER volume per unit length of dendrite did not change with LTP (control: 0.018 ± 0.001 μm^3^/μm, LTP: 0.018 ± 0.002 μm^3^/μm, hnANOVA: F_(1, 14)_ = 0.18, p = 0.67). Total synaptic input along dendritic segments was tightly correlated with total SER volume in control conditions (simple regression: r = 0.63, F_(1, 7)_ = 12.01, p < 0.05) but not with LTP (simple regression: r = 0.03, F_(1,7)_ = 0.01, p = 0.93).

SER enters less than 20% of hippocampal dendritic spines^14, 15^. In those spines that contain SER, SER exists as either a simple tubule (Fig. 3a) or as a larger and more complex spine apparatus, with folds of SER stacked between densely stained material (Fig. 3b). It is not known whether the occupancy, complexity, or volume of SER in dendritic spines changes with LTP. To explore this, SER was reconstructed in dendritic spines and identified as a simple SER tubule or as a spine apparatus. Overall, the volume of SER in spines was significantly greater during LTP, but we did not uncover a significant change in volume of either SER tubules or spine apparatuses (Fig. 3c). Under both conditions, we found similar percentages of spines containing SER (13-15%); however, there was a significant shift during LTP from tubules of SER to spines apparatuses (Fig. 3d). Thus, the increase in SER volume in spines during LTP was accounted for by the observed shift from SER tubules to larger and more complex spine apparatuses.

**Fig. 3:**
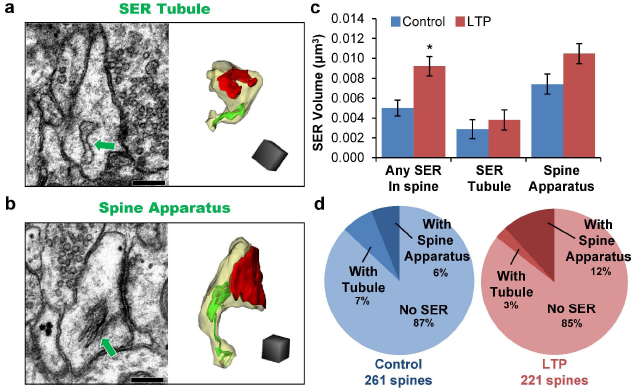
SER is more likely to form a spine apparatus during LTP. (a) Example electron micrograph and reconstruction of a dendritic spine (yellow) and PSD (red) with a single tubule of SER (green arrow in micrograph, green surface in the reconstruction). (b) Example electron micrograph and reconstruction of a dendritic spine with a spine apparatus using same color scheme as in A. Scale bars and cubes = 250 μm on each side in A and B. (c) Volume of SER in spines increased with LTP (hnANOVA: F_(1,47)_ = 6.23, p < 0.05); however, there was no difference in the volume of SER tubules in spines (F_(1, 12)_ = 1.37, p = 0.26) or in the volume of spine apparatuses during LTP (F_(1, 24)_ = 0.63, p = 0.44). (d) Percentages of spines with SER tubules, with spine apparatuses, and without SER. During LTP, there was a shift to more spines containing a spine apparatus (chi-squared test: p < 0.05).

Integral membrane proteins such as AMPARs move through the dendritic shaft more rapidly along simple, tubular SER and more slowly where SER is more complex^13^. SER tends to be more complex at the base of spines^14, 15^, which facilitates cargo delivery and ultimately receptor insertion at nearby synapses^13^. To test whether structural remodeling of dendritic SER could influence its complexity during LTP, we analyzed regions of aspiny versus spiny dendritic segments. An aspiny dendritic segment was defined as a length of dendrite at least 100 nm long without a spine origin (Fig. 4a). In the prior paper, SER complexity was estimated by a simple index of SER area summed across spiny vs. aspiny segments^13^. Here we refined this index to account for the combined effects of SER volume and branching and to normalize for dendritic segment length and caliber as defined by equation (1) (Fig. 4b, see Methods):

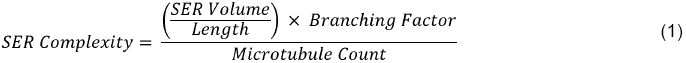

**Fig. 4:**
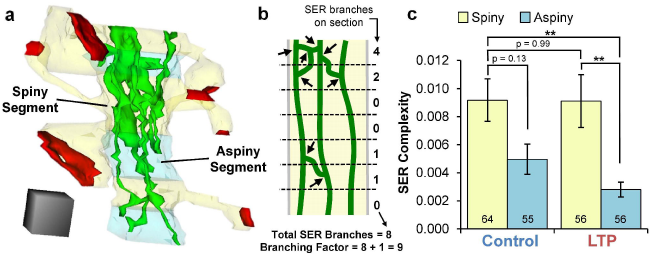
During LTP, SER is simplified in aspiny dendritic segments. (a) Example reconstruction of dendrite showing spiny (yellow) and aspiny (teal) segments, PSDs (red), and SER (green) extending through the dendritic shaft. An aspiny dendritic segment was defined as having a length of at least 100 nm without a spine origin. Scale cube = 250 μm on each side. (b) Diagram showing identification of SER (green) branches in a model dendritic segment (yellow). SER branch points (black arrows) were quantified on each EM section (separated by dotted lines) across the dendritic segment. For example, an aspiny dendritic segment 100 nm long, containing 0.0246 μm^3^ of SER with 0 branches, and 12 microtubules, would have a SER complexity value = (0.0246/.1 x (0+1))/12 = 0.0205. In contrast, a similar dendritic segment that differed only by having 1 SER branch point would have a SER complexity value = (0.0246/.1 x (1+1))/12 = 0.0410, etc. (c) During LTP, SER complexity in aspiny segments was substantially reduced relative to spiny segments in both LTP (hnANOVA: F_(3, 195)_ = 4.88, p < 0.01, Tukey post-hoc, p < 0.01) and control conditions (Tukey post-hoc, p < 0.01). In the control condition alone, the difference in SER complexity between spiny and aspiny dendritic segments did not reach statistical significance (Tukey post-hoc, p = 0.13). The SER complexity was comparable for spiny dendritic segments from both control and LTP conditions (Tukey post-hoc, p = 0.99). Sample sizes for each group are shown on the corresponding bar of the graph.

SER volume was computed for each segment by summing the SER profile areas across EM sections and multiplying by section thickness. The branching factor was computed by summing the number of branch points in each dendritic segment and then adding 1 to ensure a non-zero value for segments with unbranched tubules of SER (Fig. 4b). Microtubule count scales with dendrite caliber^21^, hence the SER complexity index was normalized by the number of microtubules to control for larger dendrites having a greater capacity for SER. Overall, SER complexity was greater in spiny versus aspiny segments of the dendrite (hierarchical nested ANOVA [hnANOVA]: F_(1, 195)_ = 12.09, p < 0.001). Furthermore, during LTP, SER complexity was sustained in spiny segments of the dendrite but substantially reduced in the aspiny segments (Fig. 4c). Thus, as spines acquired spine apparatuses during LTP (see Fig. 3d), SER was shuttled from aspiny segments to spiny segments of the dendrite. The lower complexity of SER in the aspiny segments would speed trafficking across regions of the dendrite lacking synapses.

Next, we considered whether the increase in SER in dendritic spines influenced synapse growth during LTP. Dendritic spines with polyribosomes have been shown to have larger PSDs than spines without polyribosomes at 2 hours during LTP^4, 11^. SER is also involved in protein synthesis and posttranslational modification of proteins^22^. Hence, we analyzed whether co-localization of polyribosomes and SER in spines enhanced synapse enlargement during LTP. Under both control and LTP conditions, most spines had neither polyribosomes nor SER (Fig. 5a-b, e). Some spines had either polyribosomes or SER, and a small percentage had both (Fig. 5c-d, e). Under control conditions, synapses on spines without SER were smaller than those on spines with SER, regardless of whether the spine contained a polyribosome or not (without SER: 0.057 ± 0.003 μm^2^, with SER: 0.160 ± 0.017 μm^2^, hnANOVA: F_(1,243)_ = 126.68, p < 0.001). Synapses on spines without SER showed a small but statistically significant increase in size 2 hours during LTP whether or not they contained polyribosomes (Fig. 5f). In contrast, synapses on spines with SER had a more dramatic increase in size (Fig. 5f). Interestingly, the largest synapses were on spines that contained both SER and polyribosomes during LTP. These results suggest that spines with SER were primed to undergo greater synaptic enlargement during LTP than those without SER. Such a dramatic increase in synapse size on spines containing both SER and polyribosomes suggests that these spines were able to mobilize these resources, which worked synergistically to support synapse growth during LTP.

**Fig. 5:**
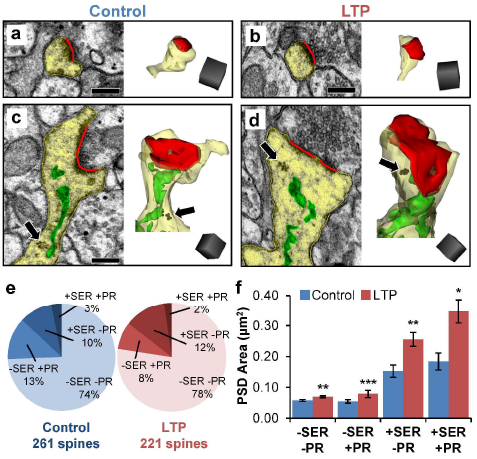
Spines with larger synapses are more likely to contain SER. (a-b) Example electron micrographs and reconstructions of dendritic spines (yellow) with no SER or polyribosomes from both conditions. (c-d) Example electron micrographs and reconstructions of dendritic spines with SER (green), PSDs (red) and polyribosomes (PR, black arrows). Scale bars = 250 μm. Scale cube = 250 μm on each side. (e) Frequency of spines with just polyribosomes (SER-PR+), just SER (SER+ PR-), neither (SER-PR-), or both (SER+ PR+) in control and LTP conditions. SER+ PR+ spines were rare and were found in control conditions of one experiment and LTP conditions in another experiment. There were no differences in the proportions of spines containing SER, PR, neither, or both between control and LTP conditions (chi-square: p = 0.35). (f) Comparison of average PSD size in control and LTP conditions on spines with or without SER or polyribosomes. Mean PSD area on spines without SER or polyribosomes was modestly larger with LTP (hnANOVA: F_a_, 347) = 8.39, p < 0.01). PSDs on spines with just polyribosomes were even larger with LTP (F(1, _35_) = 20.43, p < 0.001). PSDs on spines with just SER were larger still with LTP (F_(1, 35)_ = 9.38, p < 0.01). The largest increase in PSD area with LTP was on spines with both SER and polyribosomes (F_(1, 6)_ = 9.8167, p < 0.05).

Finally, since synapses on spines containing both SER and polyribosomes were the largest during LTP, we were interested to learn whether the complexity of SER at the base of those spines was altered. We reasoned that these spines might benefit from highly complex SER at their bases, which would serve as a local source of proteins the spine could access during synapse growth^13^. We analyzed SER complexity 0.5 μm around the base of each spine (Fig. 6a). We removed spines from this analysis if 0.5 μm around their base fell outside the length of the analyzed dendritic segment. In agreement with our findings above (see Fig. 4d), when we analyzed all spines together we found that SER complexity was retained at their bases during LTP (control: 0. 0082 ± 0.0006, LTP: 0.0077 ± 0.0008, hnANOVA: F_(1, 424)_ = 0.04, p = 0.84). Interestingly, only SER complexity at the base of spines that contained both SER and polyribosomes was significantly greater during LTP (Fig. 6b). Thus, not only is SER complexity conserved at the base of spines during LTP, SER complexity was even greater at the base of the few “privileged” spines that contained both SER and polyribosomes, spines that expanded their synapses the most during LTP. This finding suggests that LTP induces structural changes in SER that facilitate the movement of cargo to and from the largest synapses, providing a local mechanism to enhance their growth.

**Fig. 6:**
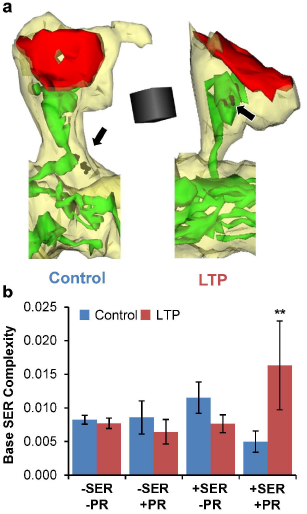
SER complexity is greatest at the base of spines that contain polyribosomes and SER during LTP. (a) Example reconstructions of dendritic spines (yellow) with PSDs (red) and SER (green) in spines containing both SER and polyribosomes from each condition. SER 0.5 μm around the base of each of the spines has also been reconstructed. (b) SER complexity 0.5 μm around the base of spines with or without SER or polyribosomes under control conditions and during LTP. During LTP, SER complexity was conserved at the base of spines without SER or polyribosomes (-SER -PR, hnANOVA: F_(1, 316)_ = 0.06, p = 0.81), at the base of spines with polyribosomes only (-SER +PR, F_(1, 30)_ = 0.13, p = 0.72), and at the base of spines with SER only (+SER -PR, F(_1, 33_) = 2.17, p = 0.15). SER complexity was, however, increased at the base of the few spines that contained both polyribosomes and SER during LTP (+SER +PR, F_(1, 5)_ = 23.15, p < 0.01).

## Discussion

Here we have demonstrated for the first time that the structure of dendritic SER is dynamic in ways that support enlargement of specific synapses during LTP in the adult hippocampus. Trafficking of membrane and proteins along SER is critical for the expression of synaptic plasticity, and movement throughout the dendrite is slowed in regions where SER structure is most complex, thereby enhancing local delivery of the cargo^13^. Under baseline conditions *in vivo*, SER is more complex in portions of the dendritic shaft with more or larger dendritic spines^13, 14^. We now show that under control conditions in adult hippocampal slices, the complexity of dendritic shaft SER was greater where total synaptic input was higher, consistent with the prior *in vivo* findings. By 2 hr during LTP, the total SER volume per dendritic segment length was unchanged, yet the structure of SER underwent substantial reorganization. Both the volume and complexity of SER increased in dendritic spines, and synapse enlargement was greatest on those spines containing SER and a polyribosome. During LTP, SER became more complex at the base of spines that contained both SER and a polyribosome, and became less complex along portions of the dendrite that lacked spines. These findings suggest that spines containing SER and polyribosomes were primed to undergo greater synapse enlargement during LTP than those lacking them. Furthermore, SER was redistributed from portions of the dendritic shaft with no spines to portions where synapses underwent the greatest enlargement during LTP.

SER also contributes to the regulation of calcium dynamics^23-27^. Elevations in calcium can be localized within spines or spread through the dendritic shaft^23, 28^, ultimately propagating to the nucleus where calcium transients influence gene transcription^29^. In the hippocampus, RyRs localize to SER in dendritic spines^30^ and respond to calcium entering through ionotropic glutamate receptors and voltage gated calcium channels^23^. RyR activation results in calcium-mediated calcium release that amplifies an otherwise weak signal^24, 31-34^. IP_3_Rs, on the other hand, are localized to SER in the dendritic shaft^25, 30^ and are activated by calcium and IP_3_^23^. Hotspots of IP_3_Rs occur along SER in CA1 dendrites where SER is more elaborate^35^. Furthermore, calcium released from SER via IP_3_R activation during LTP is involved in coordinating plasticity among synaptic sites along dendrites^36, 37^. Thus, the SER elaboration during LTP in spines and at their bases could enhance local calcium signaling and serve as a potentiating signal to sustain and enlarge those synapses^38, 39^. In contrast, where dendritic SER became simplified, less calcium would be released and phosphatases would be more likely to be activated^40, 41^, possibly leading to spine loss along these portions of the dendrite.

During LTP, the SER in dendritic spines was more likely to form a complex spine apparatus, while under control conditions spine SER usually formed just a simple tubule. Immuno-reactive markers for the Golgi apparatus, which is required for the translation and insertion of integral membrane proteins, have been identified in dendritic shafts and the spine apparatus, suggesting the spine apparatus could act as a mobile Golgi outpost^22, 42-44^. Synaptopodin is an essential component of the spine apparatus^45^ and live-imaging experiments in cultured neurons show that dendritic spines containing synaptopodin have larger AMPAR-mediated excitatory postsynaptic potentials due to ryanodine-triggered calcium release^19^. Two-photon microscopy reveals that synaptic depression is also regulated by calcium influx into large spines associated with synaptopodin^18^. Thus, spine apparatus elaboration and polyribosome recruitment to a subset of spines could serve to regulate intra-spine calcium and local protein synthesis and support enhanced bidirectional synaptic plasticity, namely enlargement during LTP, at those spines.

Several molecular mechanisms could be triggered that would link the induction of LTP to the local elaboration and redistribution of dendritic SER. One likely mechanism involves CLIMP63, an integral membrane protein in SER, and protein kinase C (PKC), which phosphorylates CLIMP63^13^ and is activated during LTP^46^. PKC-mediated phosphorylation of CLIMP63 causes SER to dissociate from microtubules and become more elaborate^13, 47, 48^. Other signaling molecules are also activated in dendritic spines and the neighboring dendritic shaft during LTP, such as calcium/calmodulin-dependent protein kinase II (CaMKII)^49^ and the small GTPase Ras^50^, which stimulates extracellular signal-regulated kinase (ERK). Together with PCK, CAMKII or ERK may also phosphorylate CLIMP63 during LTP, resulting in the elaboration of SER that would facilitate offloading of cargo and support growth of activated synapses. Further along the dendrite, away from activated spines, the dephosphorylation of CLIMP63 would cause SER to associate with microtubules and become straighter and more tubular^13^. This simplification of SER during LTP would enhance movement of proteins and other cargo away from less active synapses, possibly preventing the formation of new spines or resulting in the elimination of weak spines in those dendritic regions^4^. Thus, the dramatic reorganization of SER during LTP shown here supports an important role for SER in coordinating synaptic plasticity along adult hippocampal dendrites.

## Methods

### Physiology

All studies were done in accordance with and approved by the Institutional Animal Care and Use Committee of the University of Texas at Austin. Adult male Long-Evans rats aged 60-61 days old (319-323 g) were anesthetized with halothane and decapitated. Hippocampal slices 400 μm thick were collected from the middle third of the hippocampus (2 slices from 2 animals) and recovered in an interface chamber in artificial cerebrospinal fluid (16.4 mM NaCl, 5.4 mM KCl, 3.2 mM CaCl_2_, 1.6 mM MgSO_4_, 26.2 mM NaHCO_3_, 1.0 mM NaH_2_PO_4_, and 10 mM dextrose) for ~3 hours at 32 °C (Fig. 1A, adapted from Bourne and Harris, 2011). A recording electrode was placed in the middle of *stratum radiatum* in area CA1. Two stimulating electrodes were placed on either side of the recording electrode separated by a distance of 600-800 μm to guarantee stimulation of distinct populations of synapses^4, 11^. The initial slope of the field excitatory potential was measured and baseline recordings were collected from each stimulating electrode every 2 minutes (offset by 30 sec) for ~30 min. Theta-burst stimulation (TBS, 8 trains of 10 bursts at 5 Hz of 4 pulses at 100 Hz delivered 30 seconds apart) was delivered to one stimulating electrode at time 0 to induce LTP (Fig. 1B). The site of LTP induction (CA3 or subicular side of the recording electrode) was alternated between experiments. Responses following TBS were then monitored for 2 hours.

### Fixation and Processing for EM

The electrodes were removed and hippocampal slices were fully fixed within 1 minute of the last recording by turning the slice, still on its net, into a mixed aldehyde fixative (6% glutaraldehyde and 2% paraformaldehyde in 0.1 M cacodylate buffer with 2 mM CaCl_2_ and 4 mM MgSO_4_) and microwaving the slice in fixative for 10 sec. Slices were kept in fixative overnight at room temperature and then embedded in agarose and vibra-sliced at 70 μm (Leica WT 1000S, Leica, Nussloch, Germany). For each stimulation site, the vibra-slice containing the electrode indentation along with two adjacent vibra-slices were processed for EM through a 1% osmium/1.5% potassium ferrocyanide mixture, 1% osmium alone, dehydrated through graded ethanols (50-100%) and propylene oxide, embedded in LX112, and placed in a 60 °C oven for 48 hours^4^. Approximately 200 serial sections were collected 150-200 μm lateral to each electrode at a depth of 120-150 μm from the air surface of the slice and were mounted on pioloform-coated slot grids (Synaptek, Ted Pella Inc., Redding, CA). Sections were counterstained with ethanolic uranyl acetate and Reynolds lead citrate. The serial sections and a calibration grid (Ted Pella Inc.) were then imaged on a JEOL 1230 transmission electron microscope (Peabody, MA) with a Gatan digital camera.

### Three-dimensional Reconstructions

Serial section images were coded so as to remain blind to condition and were imported into RECONSTRUCT™ (freely available at synapses.clm.utexas.edu) and aligned. Section thickness was computed using the cylindrical diameters method by dividing the diameters of longitudinally sectioned mitochondria by the number of sections the mitochondria spanned^4^. In this study, SER was traced in dendrites that spanned at least 100 serial sections and had 9-22 microtubules, a measure of dendrite caliber^4, 21^.

### Identification of SER Branches

To identify points where SER branched, we wrote a simple script in Python, which treated individual SER traces created in RECONSTRUCT™ as vertices. Edges between vertices were identified when the traces overlapped one another on adjacent serial sections. We defined the number of SER branching events existing on a particular EM section, *b*(*v*), as in equation (2):

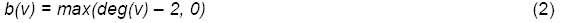

The degree of a vertex v, *deg*(*v*), was the number of edges to which the vertex belonged. Tubular, nonbranching SER traces existed when the vertex had no neighboring traces (0 edges, *deg*(*v*) = 0), had one neighboring trace (1 edge, *deg*(*v*) = 1), or had two neighboring traces (2 edges, *deg*(*v*) = 2). Branching SER existed when the vertex had > 2 neighbors (*deg*(*v*) > 2). This metric proceeded linearly.

### Statistical Analyses

In this study, 9 control dendrites (4 from animal 1, 5 from animal 2) and 9 LTP dendrites (5 from animal 1, 4 from animal 2) were analyzed. Statistical analyses were performed in R (r-project.org) and STATISTICA (StatSoft, Tulsa, OK). Hierarchical nested ANOVAs (hnANOVAs) were used (with dendrite nested in condition and experiment and experiment nested in condition) to ensure results were not driven by a particular dendrite or experiment. The sample sizes of spines with SER and polyribosomes versus those without varied widely. Therefore, for LTP-related comparisons of data categorized by spine content (SER and/or polyribosomes), we performed separate hnANOVAs on each group. The sample sizes of dendritic segments categorized as spiny or aspiny were comparable, thus a hnANOVA was performed across these groups followed by Tukey’s HSD post-hoc test to determine significant differences among the groups. Simple regression was used to investigate the effect of a continuous predictor on a dependent variable (e.g., PSD area v. SER volume) and chi-square tests were used to investigate changes in proportions of spines categorized by content during LTP. Statistical tests are reported in results and figure legends where appropriate. Significance was set to p < 0.05 and asterisks in figures denote p < 0.05 (*), p < 0.01 (**), and p < 0.001 (***).

## Acknowledgments

This study was supported by NIH Grants NS21184, MH095980, and NS074644 to KMH, NS71442 to JNB, and the Texas Emerging Technologies Fund. JNB and KMH designed the experiments. JNB performed the experiments. LFL wrote the SER branching script in Python. MAC and JNB analyzed the data. MAC, JNB, and KMH wrote the manuscript. We thank Dr. Daniel Johnston, Dr. Deborah Watson, and Dr. Maria E. Bell, for helpful discussions concerning this study.

## References

1. Bosch, M. & Hayashi, Y. Structural plasticity of dendritic spines. Current Opinion in Neurobiology 22, 383–388 (2012).

2. Meyer, D., Bonhoeffer, T. & Scheuss, V. Balance and stability of synaptic structures during synaptic plasticity. Neuron 82, 430–443 (2014).

3. Yuste, R. & Bonhoeffer, T. Morphological changes in dendritic spines associated with long-term synaptic plasticity. Annu. Rev. Neurosci. 24, 1071–1089 (2001).

4. Bourne, J. N. & Harris, K. M. Coordination of size and number of excitatory and inhibitory synapses results in a balanced structural plasticity along mature hippocampal CA1 dendrites during LTP. Hippocampus 21, 354–373 (2011).

5. Geinisman, Y., deToledo-Morrell, L. & Morrell, F. Induction of long-term potentiation is associated with an increase in the number of axospinous synapses with segmented postsynaptic densities. Brain Res. 566, 77–88 (1991).

6. Ehlers, M. D. Dendritic trafficking for neuronal growth and plasticity. Biochem. Soc. Trans. 41, 1365–1382 (2013).

7. Park, M. et al. Plasticity-induced growth of dendritic spines by exocytic trafficking from recycling endosomes. Neuron 52, 817–830 (2006).

8. Isaac, J. T., Nicoll, R. A. & Malenka, R. C. Evidence for silent synapses: implications for the expression of LTP. Neuron 15, 427–434 (1995).

9. Park, M., Penick, E. C., Edwards, J. G., Kauer, J. A. & Ehlers, M. D. Recycling endosomes supply AMPA receptors for LTP. Science 305, 1972–1975 (2004).

10. Bourne, J. N., Sorra, K. E., Hurlburt, J. & Harris, K. M. Polyribosomes are increased in spines of CA1 dendrites 2 h after the induction of LTP in mature rat hippocampal slices. Hippocampus 17, 1–4 (2007).

11. Ostroff, L. E., Fiala, J. C., Allwardt, B. & Harris, K. M. Polyribosomes redistribute from dendritic shafts into spines with enlarged synapses during LTP in developing rat hippocampal slices. Neuron 35, 535–545 (2002).

12. Higley, M. J. & Sabatini, B. L. Calcium signaling in dendrites and spines: practical and functional considerations. Neuron 59, 902–913 (2008).

13. Cui-Wang, T. et al. Local zones of endoplasmic reticulum complexity confine cargo in neuronal dendrites. Cell 148, 309–321 (2012).

14. Spacek, J. & Harris, K. M. Three-dimensional organization of smooth endoplasmic reticulum in hippocampal CA1 dendrites and dendritic spines of the immature and mature rat. J Neurosci 17, 190–203 (1997).

15. Cooney, J. R., Hurlburt, J. L., Selig, D. K., Harris, K. M. & Fiala, J. C. Endosomal compartments serve multiple hippocampal dendritic spines from a widespread rather than a local store of recycling membrane. Journal of Neuroscience 22, 2215–2224 (2002).

16. Deller, T., Merten, T., Roth, S. U., Mundel, P. & Frotscher, M. Actin-associated protein synaptopodin in the rat hippocampal formation: localization in the spine neck and close association with the spine apparatus of principal neurons. J Comp Neurol 418, 164–181 (2000).

17. Fifková, E., Markham, J. A. & Delay, R. J. Calcium in the spine apparatus of dendritic spines in the dentate molecular layer. Brain Res. 266, 163–168 (1983).

18. Holbro, N., Grunditz, A. & Oertner, T. G. Differential distribution of endoplasmic reticulum controls metabotropic signaling and plasticity at hippocampal synapses. Proc Natl Acad Sci USA 106, 15055–15060 (2009).

19. Vlachos, A. et al. Synaptopodin regulates plasticity of dendritic spines in hippocampal neurons. Journal of Neuroscience 29, 1017–1033 (2009).

20. Korkotian, E., Frotscher, M. & Segal, M. Synaptopodin regulates spine plasticity: mediation by calcium stores. Journal of Neuroscience 34, 11641–11651 (2014).

21. Fiala, J. C. et al. Timing of neuronal and glial ultrastructure disruption during brain slice preparation and recovery in vitro. J Comp Neurol 465, 90–103 (2003).

22. Pierce, J. P., Mayer, T. & McCarthy, J. B. Evidence for a satellite secretory pathway in neuronal dendritic spines. Curr Biol 11, 351–355 (2001).

23. Berridge, M. J. Neuronal calcium signaling. Neuron 21, 13–26 (1998).

24. Raymond, C. R. & Redman, S. J. Different calcium sources are narrowly tuned to the induction of different forms of LTP. Journal of Neurophysiology 88, 249–255 (2002).

25. Simpson, P. B., Challiss, R. A. & Nahorski, S. R. Neuronal Ca2+ stores: activation and function. Trends in Neurosciences 18, 299–306 (1995).

26. Emptage, N., Bliss, T. V. & Fine, A. Single synaptic events evoke NMDA receptor-mediated release of calcium from internal stores in hippocampal dendritic spines. Neuron 22, 115–124 (1999).

27. Sala, C., Roussignol, G., Meldolesi, J. & Fagni, L. Key role of the postsynaptic density scaffold proteins Shank and Homer in the functional architecture of Ca2+ homeostasis at dendritic spines in hippocampal neurons. Journal of Neuroscience 25, 4587–4592 (2005).

28. Segal, M. & Korkotian, E. Endoplasmic reticulum calcium stores in dendritic spines. Front Neuroanat 8, 64 (2014).

29. Bading, H. Nuclear calcium signalling in the regulation of brain function. Nat Rev Neurosci 14, 593–608 (2013).

30. Sharp, A. H. et al. Differential immunohistochemical localization of inositol 1,4,5-trisphosphate-and ryanodine-sensitive Ca2+ release channels in rat brain. J Neurosci 13, 3051–3063 (1993).

31. Verkhratsky, A. & Shmigol, A. Calcium-induced calcium release in neurones. Cell Calcium 19, 1–14 (1996).

32. Raymond, C. R. & Redman, S. J. Spatial segregation of neuronal calcium signals encodes different forms of LTP in rat hippocampus. The Journal of Physiology 570, 97–111 (2006).

33. Blaustein, M. P. & Golovina, V. A. Structural complexity and functional diversity of endoplasmic reticulum Ca(2+) stores. Trends in Neurosciences 24, 602–608 (2001).

34. Rose, C. R. & Konnerth, A. Stores not just for storage. intracellular calcium release and synaptic plasticity. Neuron 31, 519–522 (2001).

35. Fitzpatrick, J. S. et al. Inositol-1,4,5-trisphosphate receptor-mediated Ca2+ waves in pyramidal neuron dendrites propagate through hot spots and cold spots. The Journal of Physiology 587, 1439–1459 (2009).

36. Nagase, T. et al. Long-term potentiation and long-term depression in hippocampal CA1 neurons of mice lacking the IP(3) type 1 receptor. Neuroscience 117, 821–830 (2003).

37. Nishiyama, M., Hong, K., Mikoshiba, K., Poo, M. M. & Kato, K. Calcium stores regulate the polarity and input specificity of synaptic modification. Nature 408, 584–588 (2000).

38. Sajikumar, S., Li, Q., Abraham, W. C. & Xiao, Z. C. Priming of short-term potentiation and synaptic tagging/capture mechanisms by ryanodine receptor activation in rat hippocampal CA1. Learn. Mem. 16, 178–186 (2009).

39. Mellentin, C., Jahnsen, H. & Abraham, W. C. Priming of long-term potentiation mediated by ryanodine receptor activation in rat hippocampal slices. Neuropharmacology 52, 118–125 (2007).

40. Lisman, J. A mechanism for the Hebb and the anti-Hebb processes underlying learning and memory. Proc Natl Acad Sci USA 86, 9574–9578 (1989).

41. Mulkey, R. M., Herron, C. E. & Malenka, R. C. An essential role for protein phosphatases in hippocampal long-term depression. Science 261, 1051–1055 (1993).

42. Gardiol, A., Racca, C. & Triller, A. Dendritic and postsynaptic protein synthetic machinery. J Neurosci 19, 168–179 (1999).

43. Grigston, J. C., VanDongen, H. M. A., McNamara, J. O. & VanDongen, A. M. J. Translation of an integral membrane protein in distal dendrites of hippocampal neurons. Eur J Neurosci 21, 1457–1468 (2005).

44. Horton, A. C. et al. Polarized secretory trafficking directs cargo for asymmetric dendrite growth and morphogenesis. Neuron 48, 757–771 (2005).

45. Deller, T. et al. Synaptopodin-deficient mice lack a spine apparatus and show deficits in synaptic plasticity. Proc Natl Acad Sci USA 100, 10494–10499 (2003).

46. Malinow, R., Schulman, H. & Tsien, R. W. Inhibition of postsynaptic PKC or CaMKII blocks induction but not expression of LTP. Science 245, 862–866 (1989).

47. Vedrenne, C., Klopfenstein, D. R. & Hauri, H.-P. Phosphorylation controls CLIMP-63-mediated anchoring of the endoplasmic reticulum to microtubules. Mol. Biol. Cell 16, 1928–1937 (2005).

48. Klopfenstein, D. R., Kappeler, F. & Hauri, H. P. A novel direct interaction of endoplasmic reticulum with microtubules. EMBO J. 17, 6168–6177 (1998).

49. Lee, S.-J. R., Escobedo-Lozoya, Y., Szatmari, E. M. & Yasuda, R. Activation of CaMKII in single dendritic spines during long-term potentiation. Nature 458, 299–304 (2009).

50. Harvey, C. D., Yasuda, R., Zhong, H. & Svoboda, K. The spread of Ras activity triggered by activation of a single dendritic spine. Science 321, 136–140 (2008).

